# High-throughput, multiplex microfluidic test strip for the determination of antibiotic susceptibility in uropathogenic *E. coli* with smartphone detection

**DOI:** 10.1101/2021.05.28.446184

**Authors:** Sarah H. Needs, Zara Rafaque, Wajiha Imtiaz, Partha Ray, Simon Andrews, Alexander D. Edwards

## Abstract

Antibiotic resistance in urinary tract infections is a major global challenge and improved cost-effective and high throughput antibiotic susceptibility tests (AST) are urgently needed to inform correct antibiotic selection. We evaluated a high throughput microfluidic test strip for AST and minimum inhibitory concentration (MIC) determination in 20 urinary pathogenic *E. coli* (UPEC) isolates using six commonly prescribed or therapeutically beneficial antibiotics. The microfluidic MIC performs broth microdilution in 1 microliter volume capillaries, 100 X smaller than standard broth microdilution. Each test strip contains 10 parallel capillaries which are dipped into a single well of a 96 well plate, significantly increasing throughput over a microtitre plate. When tested with clinical UPEC isolates at standardised inoculum density, these devices gave 100% essential agreement (+/- 1 doubling dilution of antibiotic) to the gold standard microplate broth microdilution method described by CLSI. Although for some antibiotic/isolate combinations an earlier endpoint readout reduced accuracy, MIC test strips read at a 6h endpoint still gave 69 – 100 % essential agreement depending on the antibiotic. Growth could be detected significantly earlier than 6h, but with a trade-off between speed vs accuracy. These high-throughput, multiplexed test strips could be used to increase throughput and give faster results than microplates while retaining the core broth microdilution methodology of gold standard techniques for AST and MIC determination.

## 1 Introduction

Although improved diagnostic technology is widely acknowledged to be essential to combat antimicrobial resistance (AMR) (Trevas et al., 2020), antibiotic susceptibility testing (AST) of clinical samples such as urine remains challenging. Although urinary tract infections (UTI) are often self-limiting, a high prevalence places a significant burden on health systems, and when infections escalate hospital treatment can be required. Current primary care and hospital treatment protocols rely on empirical antibiotic administration at first presentation (Grigoryan et al., 2014, O’Grady et al., 2019), which exacerbates the clinical and economic burden because of a high level of broad and rapidly changing AMR profile necessitating constant updates to guidelines. When urine samples are taken and tested, adjustment of antibiotic is often required when results are returned days later. Resistance ranges up to ~50% of uropathogenic *E. coli* (UPEC) isolates in OECD countries and as high as >80% outside OECD, depending on antibiotic (Bryce et al., 2016) which limit the use of many common antibiotics, and reduce effectiveness of empirical prescribing. In antibiotics with resistance rates > 20% empiric treatment is unsuitable (O’Grady et al., 2019). This leads to consequences for the patient (infection may not be controlled if pathogen is resistant), the clinician (limited treatment choices) and the public (emergence and spread of AMR and subsequent drug-resistant infections). Even if susceptibility testing is requested, urine culture plus AST takes 2-3 days, during which time infections worsen if the empirical antibiotic is ineffective, or if treatment is delayed prior to culture results.

Several emerging strain-characterisation technologies to determine AST have been developed to the point of large scale uptake into central microbiology laboratories (Pulido et al., 2013). Many labs are now using mass spectroscopy for identification of clinically important microbes by peptide mass fingerprinting to replace traditional analytical microbiology techniques (Lavigne et al., 2013, Florio et al., 2020). Nucleic acid tests (NAT) such as PCR can detect specific AMR genes, allowing rapid identification of known resistance markers, but do not always reliably predict phenotypic antibiotic susceptibility and can only detect target sequences – as these targets change, tests will need regular updating informed by phenotypic-genotypic surveillance programs (Needs et al., 2020). Phenotypic AST, including the current gold standard methods (Andrews, 2001, Andrews, 2005, Jorgensen and Turnidge, 2015) of broth microdilution (BMD) performed in a microtitre plate (MTP) and disc diffusion assays performed on agar plates, demonstrate antibiotic suitability by direct phenotypic identification of which antibiotics achieve bactericidal or bacteriostatic effects (i.e. susceptible), vs which isolates grow in the presence of antibiotic (i.e. resistant). Phenotypic AST currently requires a pure bacterial culture isolated by overnight plating, laborious handling and requiring centralised laboratory equipment. There are several automated phenotypic AST machines currently available. The Vitek^®^ 2 (Biomerieux) reports bacterial identification and MIC results within 5-8 h, following bacterial colony isolation after overnight agar plating. In contrast, the Accelerate Pheno^R^ system is designed to be used directly with samples (Kaprou et al., 2021). These methods are able to reduce the time to result by hours and potentially days (Kinn et al., 2019) but still require the use of centralised labs for testing.

Microfluidic technologies offer several advantages, especially in point-of-care applications, including portability and rapid result times. Recently many examples of microfluidic analytical microbiology devices illustrate the potential for AST miniaturisation (Avesar et al., 2017, Azizi et al., 2018, Wu and Dekker, 2016, Kim et al., 2015, Kang et al., 2019). A “millifluidic” system made from simple FEP tubing illustrated that analysis of AST can be miniaturised using microdroplets allowing parallel analysis of large populations of individual cells (Baraban et al., 2011). Whilst this microdroplet system offers improved resolution in quantitative AST (Jiang et al., 2016) pure cultures from overnight plating and complex instrumentation are still required. Microfluidic devices have also been developed for quantitative susceptibility testing. Multi-chamber devices can distribute bacterial samples into chambers loaded with antibiotics (Cira et al., 2012, Avesar et al., 2017, Azizi et al., 2018, He et al., 2020, Matsumoto et al., 2016), or cells can be observed in microfluidic systems that generate antibiotic gradients (Kim et al., 2019, Li et al., 2014). Microfluidic culture can be combined with digital microscopy and image analysis to allow direct observation of the effects of antibiotics on cell behaviour or growth (Choi et al., 2013, Wistrand-Yuen et al., 2020, Baltekin et al., 2017, Matsumoto et al., 2016). However, microscopic observation of bacterial growth inhibition by antibiotics does not directly map to traditional BMD or disc diffusion results, and extensive equivalency testing is required. Electrochemical detection has been combined with the conventional colorimetric metabolic dye resazurin to detect bacterial growth in microfluidic devices (Besant et al., 2015). Most microfluidic microbiology studies have focused either on exploring microdevice engineering science, or on addressing fundamental biology research questions. One notable exception is a microplate-based microfluidic AST device that has been developed into a fully automated product for testing positive blood cultures to rapidly select antibiotics for the treatment of sepsis (Choi et al., 2014, Choi et al., 2013). However, this laboratory device still requires pre-incubation of blood sample prior to AST, to achieve a threshold level of organisms required for testing.

Incubation time and inoculum size are critical parameters in AST and deviation from standardised methods can lead to errors in categorical agreement compared to the gold standard. It is therefore important to understand how both of these may affect any new AST devices or methods, however there are limited data in the literature exploring how these parameters affect microfluidic analytical microbiology.

Shorter incubation times of 6-8 h have been explored for disc diffusion assays (Mancini et al., 2020, Hombach et al., 2017, Fröding et al., 2017, Chandrasekaran et al., 2018). While categorical agreement differed depending on antibiotic/bacteria tested, overall agreement was generally >80% and it was concluded this approach was useful as a preliminary result to base urgent clinical decisions on (Fröding et al., 2017). Disc diffusion assays of 11123 Enterobacteroles/antibiotic combinations incubated for 6 h found sharper, more defined areas of inhibition with categorical agreement of 89.8% and major errors and very major errors below 4%; after 8 h the categorical agreement rose to 98.5%. The most common errors came from strains categorised as intermediate (Mancini et al., 2020); for such strains where the inhibitory concentration for antimicrobial is close to the breakpoint for scoring resistance vs susceptibility, there is always greater uncertainty in AST, even with gold standard methods. In a clinical setting it is important to identify resistant strains early, and false susceptibility is more of a risk than false resistance as the former can lead to treatment failure, so for rapid test methods to be clinically useful must be optimised to minimise false susceptibility.

The inoculum cell density is likewise an important factor when measuring MIC, and an increased starting inoculum can increase the observed MIC for a number of reasons. This well-known phenomenon is termed the inoculum effect (IE), and varying the starting bacterial cell density can influence inhibition of growth by many antimicrobials; hence guidelines for broth microdilution define acceptable ranges for inoculum density. Even within the CLSI recommended range of inoculum density, categorical errors in antibiotic sensitivity have been observed. An 8-fold difference was found in MIC between 2-8×10^5^ CFU/mL for carbapenem-resistant *Enterobacteriaceae* with meropenem (Smith and Kirby, 2018). Estimating and adjusting cell density therefore remains an essential but laborious step for many functional microbiology methods.

Healthcare budgets remain a major constraint, with high prevalence of UTI resulting in very large volume of patients presenting with UTI symptoms in primary care. AMR is more prevalent in parts of the world with most limited healthcare budgets (Pokharel et al., 2019). Very low device costs are therefore needed and expensive capital equipment should be avoided, and highly skilled technical sample processing or complex interpretation of results are likewise undesirable. There remains an urgent need for novel AST methods that can reduce workload, provide high quality data and directly test patient urine to without the need for isolation or overnight plating. Ideally new methods would replicate conventional phenotypic assays as closely as possible, to permit ease of interpretation by clinicians accustomed to current gold standard methods. For example, miniaturised BMD utilising growth detection that is similar to current automated lab BMD equipment would accelerate adoption of new technology by permitting simple clinical interpretation of similar results measured and presented in similar ways to current gold standard AST, such as MIC values and scoring resistance/susceptibility at the same breakpoint concentrations. As breakpoint concentrations are constantly monitored, ensuring new methods can make direct use of current guidelines is vital to avoid continuous equivalency testing.

Here, we fully evaluate a high throughput system for analytical microbiology that uses a fluoropolymer microcapillary film (MCF) that contains a parallel array of ten microcapillaries. The MCF is manufactured by melt-extrusion and multiplex test strips can be produced in large batches (Barbosa et al., 2015, Castanheira et al., 2015, Barbosa et al., 2014, Edwards et al., 2011), allowing us to make and test hundreds of very low-cost test devices, representing thousands of individual microcapillary tests. These devices incorporate a hydrophilic polymer coating to modify the internal walls of the capillaries allowing sample uptake by capillary action. Using this method, samples can be loaded easily by dipping the test strips into the well of a 96-well plate with the sample drawn up into the 10 capillaries by capillary action. Previously, we showed that these “Lab-on-a-stick” devices can perform a range of classical microbiology tests to identify and count viable cells (Reis et al., 2016, Needs et al., 2019, Dönmez et al., 2020, Needs et al., 2021). Here we present the first full demonstration of MIC measurement and antibiotic susceptibility testing for a range of antibiotics in clinically relevant UPEC isolates. We explore the impact of inoculum cell density on microfluidic broth microdilution and assess the kinetics of antibiotic susceptibility testing in a convenient miniaturised format, to understand better how to use microfluidics to speed up analytical microbiology.

## 2 Material and Methods

### 2.1 Bacterial Isolates

Reference strain *E. coli* 25922 was purchased from ATCC and was used as a quality control. *E. coli* 13352 was purchased from NCTC. The uropathogenic *E. coli* (UPEC) strains were collected at a tertiary care hospital of Pakistan from community acquired UTI patients (Rafaque et al., 2018, Rafaque et al., 2019) under a study that was approved by the Ethical Review Board (ERB) of the Pakistan Institute of Medical Sciences. UPEC were identified using standard microbiological and biochemical tests as described before (Ali et al., 2019).

### 2.2 Preparation of antibiotic loaded microcapillary test strips

MCF of 1 to 5 m lengths were given an internal hydrophilic coating by incubation with a 5 mg/mL solution of polyvinyl alcohol (PVOH) in water (MW 146,000-186,000, >99% hydrolysed, Sigma-Aldrich, UK) at room temperature for 2h (Pivetal et al., 2017, Reis et al., 2016). Coated strips were washed with 5 ml of PBS with 0.5 % Tween 20 (Sigma-Aldrich, UK) to remove residual PVOH, and dried on a vacuum manifold for 20 minutes per metre using a SLS Lab Basics Mini Vacuum Pump with PTFE Coated Diaphragm (Scientific Laboratory Supplies, UK).

Ciprofloxacin, cefoxitin, trimethoprim and amoxicillin were purchased from Sigma Aldrich. Nitrofurantoin and Cephalexin were from purchased from Fischer Scientific. Serial dilutions of antibiotics in sterilised milliQ water were injected using a 30-gauge needle into individual capillaries in up to 1.5 m long strips of MCF. The MCF strip was cut to 17 mm individual test strips and frozen overnight at – 80 °C. Test strips were freeze-dried for >4 h on an Edwards Modulyo freeze drier. Test strips were vacuum packed and stored at −20 °C until use.

### 2.3 Microplate Broth Microdilution

Bacterial inocula were prepared according to CLSI standard for growth preparation. Briefly, 3-5 individual colonies were inoculated into Mueller-Hinton broth and grown for 4-6 h until visibly turbid. The suspension was diluted to 0.5 McFarland equivalent which corresponds to approximately 1×10^8^ CFU/mL. The culture was diluted 1:150 in Mueller-Hinton to give 1×10^6^ CFU/mL suspension and finally 50μL was added to 50μL of antibiotic solutions in a microplate to give a 1:2 dilution to give a final suspension of bacteria at 5×10^5^ CFU/mL in two-fold dilutions of antibiotic. For BMD including resazurin, dye was added to a final concentration of 0.25 mg/mL. Plates were incubated at 37 °C overnight. MIC measurements were measured in duplicate (Fig 1a). The MIC was recorded as the lowest concentration of antibiotic that did not show resazurin conversion or turbidity. For tests that varied within duplicate measurements, the highest MIC was recorded. A growth control (no antibiotic) and a sterile control (MH broth only) was included for all isolates.

**Figure 1.**
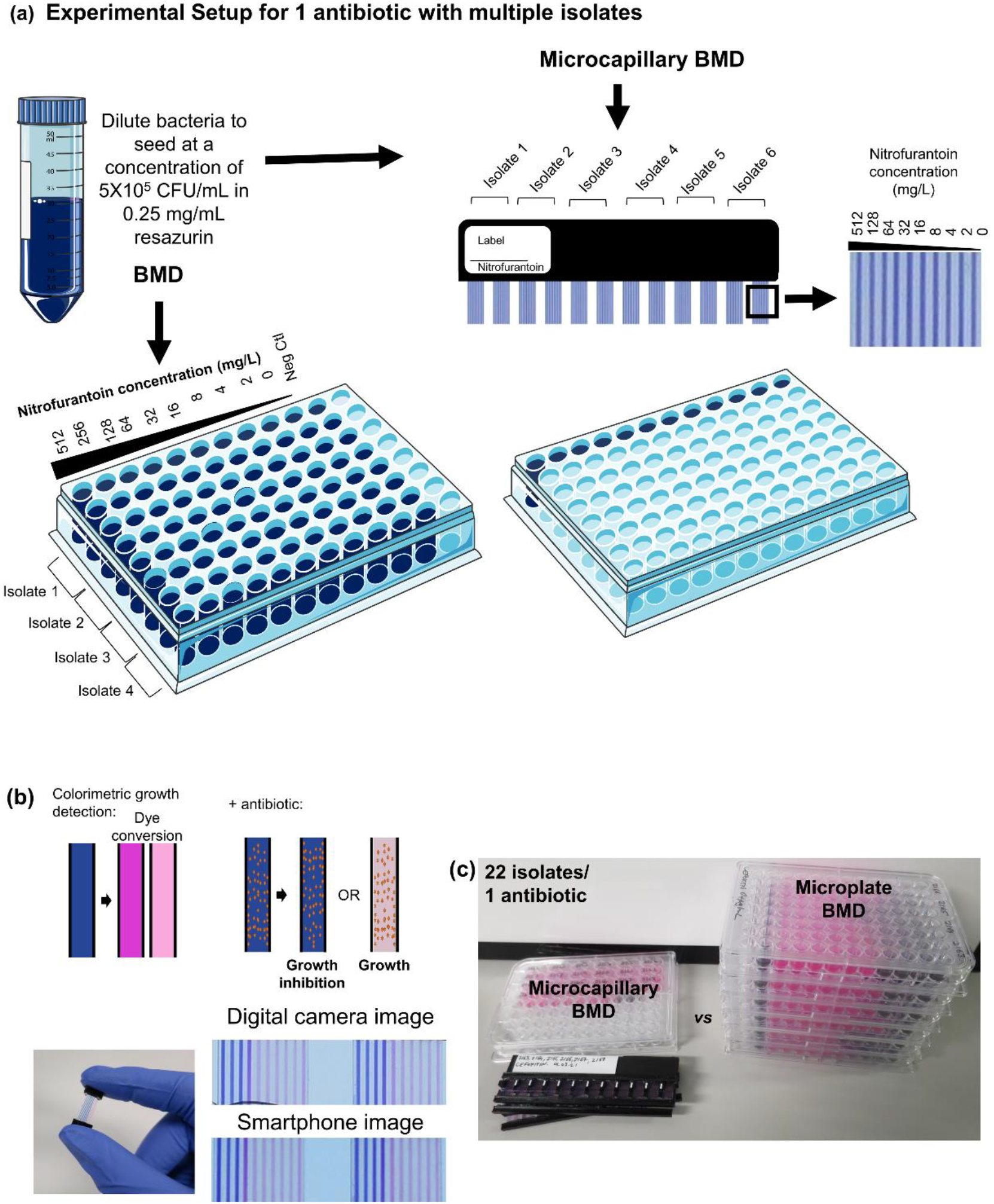
Microcapillary test use. **(a)** An array of up to 12 test strips were clipped into a ‘ladder’ holder with a 9mm pitch. Each test strip contains 10 capillaries each loaded to release doubling dilutions of antibiotic. Test strip arrays were dipped directly into 96-well microtitre plate wells allowing 1 microlitre of the sample to be drawn up by capillary action into all 10 microcapillaries. Plastic end covers filled with silicone grease were slid over the ends of the test strips, sealing the capillary ends to prevent evaporation. Image created using servier medical art **(b)** Bacteria growth and antibiotic inhibition is determined by resazurin color change. Blue to pink/white indicates growth and if antibiotic is present, indicates that concentration is below the minimum inhibitory concentration (MIC). At or above MIC, the capillary remains blue. **(c)** Each ladder of microcapillary BMD test strips provides up to 16X higher throughput than microtitre plates. Image illustrates the number of plates vs microcapillaries required to test 22 isolates in duplicate for a single antibiotic at 9 concentrations plus no antibiotic.

### 2.4 Microcapillary Broth Microdilution

Individual 17 mm test strips were clipped into 3D printed holders, able to hold up to 12 test strips. For MIC tests in MCF strips, bacteria inoculum was prepared as per the CLSI standard for growth preparation as before at a final concentration of 5×10^5^ CFU/mL and mixed in a 96 well plate with a final concentration of 0.25 mg/mL resazurin in a total volume of 200 μL. The test strips were dipped into each well. Once all capillaries were filled (approximately 3 seconds) end covers filled with silicone grease were placed on each end to stop sample evaporation. Samples were incubated overnight at 37 °C and color change monitored using an in-house automated raspberry pi camera imaging system (Needs et al., 2019). MIC measurements were taken in duplicate with two wells used for each measurement (Fig 1a). The MIC was recorded as the lowest concentration of antibiotic that did not show resazurin conversion or turbidity. For tests that varied in duplicate measurements, the highest MIC was recorded. A growth control (no antibiotic) capillary was included for all test strips.

For microcapillary BMD experiments using a higher inoculum density, CFU/mL was calculated using a spot plating protocol. Briefly, 10 μL of serially diluted culture were taken from the microtitre wells and plated on both LB agar and the Gram-negative selective media, MacConkey agar to determine cell count and ensure no contamination with other organisms. Plates were incubated overnight at 37 °C.

### 2.5 Data Analysis

Two recordings of the test strips were made. A single endpoint was recorded by colorimetric measurement after overnight incubation at 37 °C using a smartphone camera (iPhone 6s) or digital camera (Canon Powershot S120). The strips were placed on an even white light illumination screen. Isolates were scored suspectable or resistant based on EUCAST v11.0 breakpoint values in Table 1.

**Table 1.** EUCAST v 11.0 antibiotic MIC breakpoints used for categorical analysis.

| Antibiotic | EUCAST MIC breakpoints (mg/L) |
| --- | --- |
|  | ≤S/>R |
| Cefoxitin | ≤8 |
| Ciprofloxacin | ≤0.25 |
| Trimethoprim | ≤4 |
| Nitrofurantoin | ≤64 |
| Cephalexin | ≤16 |
| Amoxicillin | ≤8 |

For time-lapse imaging, resazurin conversion was recorded every 15 minutes using the POLIR robot (Needs et al., 2019), with 3280 x 2464 resolution images taken with a Raspberry Pi v2 camera. MatLab scripts were used to analyse time-lapse image series of bacterial growth in MCF, and the code can be accessed here: https://gitlab.com/sneeds/code-repository. Briefly, color images were split into red, blue and green (RGB) channels and the red channel analysed for absorbance using:

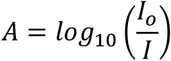

Time to resazurin conversion was calculated at which half the starting absorbance in the red channel was reached.

## 3 Results

### 3.1 Broth microdilution in microcapillary test strips accurately determines MIC for uropathogenic *E. coli*

The novel microcapillary BMD method conducts AST in dip-and-test microcapillary film (MCF) strips, with each test strip containing 10 different conditions (9 antibiotic concentrations plus no antibiotic control) in 1 μL samples, thereby reducing the test volume over 100-fold from microplates. To understand if miniaturisation affects BMD performance, we validated this microcapillary BMD AST, by comparing to a standard microplate BMD following CLSI guidelines. Growth was quantified using the colorimetric growth indicator, resazurin, and although full growth kinetics were recorded, we initially focussed on an overnight endpoint readout. Microplate BMD and microcapillary BMD were performed in parallel, testing a panel of 6 antibiotics commonly used in the treatment of UTIs, against an *E. coli* quality control (QC) strain, an *E. coli* reference strain, and 20 UPEC isolates.

*E. coli* cultured in Mueller-Hinton broth were diluted to a standard inoculum cell density of 5×10^5^ CFU/mL in Mueller-Hinton broth with a final resazurin concentration of 0.25 mg/mL in both microplates and MCF test strips in parallel, and overnight colorimetric endpoint images taken to score growth/no growth, and the same colour changes were observed in both microplate and microcapillaries (Fig 2). The capillaries in MCF test strips have a 20-fold shorter light pathlength than a microtitre plate, therefore the concentration of resazurin had to be high enough to be clearly visible in the capillaries. Endpoint colour change was scored after overnight growth from digital images, with pink/white turbid capillaries indicating bacterial growth whereas samples with a susceptible bacterial population remained blue (Fig 1b and 2).

**Figure 2.**
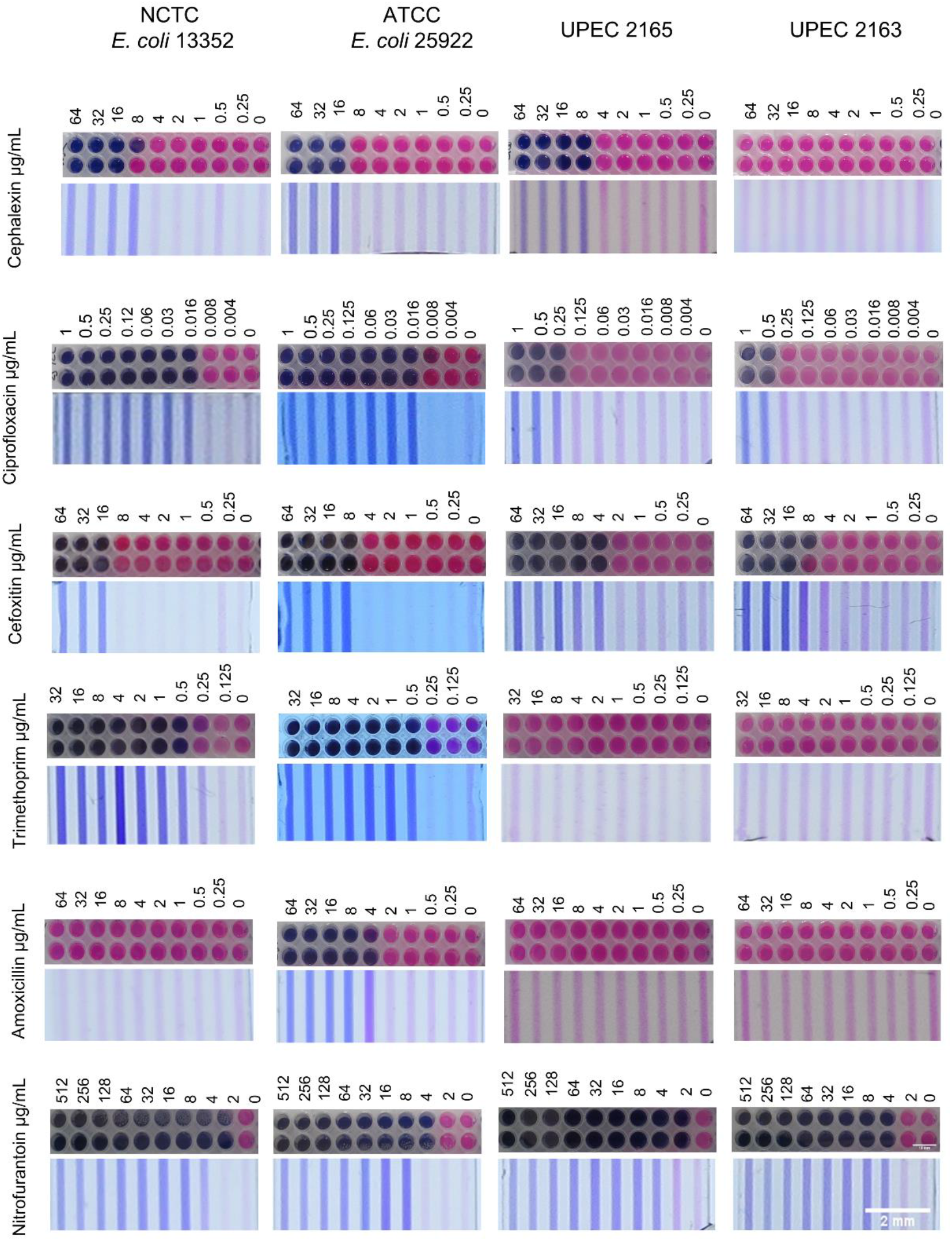
Agreement between microwell broth microdilution and microcapillary film MIC. Representative images showing MIC determination of two reference strains, *E. coli* 25922 and *E. coli* 13352 and urinary pathogen *E. coli* isolate 2165 MIC in microplate and microcapillary film. Error bar on microplates indicate 10 mm. Error bar on microcapillary film indicates 2 mm. Similar agreement was seen with all isolates tested in duplicate.

BMD in MCF and microplates was performed in duplicate and the highest MIC value for each isolate/antibiotic combination was recorded in Table S1. A total of 2640 data points were collected from the combination of 6 antibiotics and 22 samples (20 UPEC isolates plus 1 QC strain and 1 reference strain, totalling 132 antibiotic/isolate combinations). The *E. coli* 25922 strain was used as the QC strain. All MIC measured by microplate BMD and microcapillary BMD were within the acceptable MIC range according to EUCAST QC Table v 11.0 for this QC strain. Out of all combinations of *E. coli* strains tested against the 6 antibiotics, there were 56 combinations where MIC was measurable and for these there was 100% (56/56) essential agreement between microcapillary and microplate, with MIC values for each method within ±1 doubling dilution of antibiotic. No MIC value was attributed to 74 microplate BMD tests and 71 microcapillary BMD test strips. Of these, no MIC value was attributed for 3 microplate BMD conditions in which growth was only detected in the control without antibiotic (i.e. MIC was below the lowest concentration; scored as susceptible), and for 71 microplate BMD and 71 microcapillary BMD test strips where all antibiotic containing capillaries showed growth thus MIC was greater than the highest included concentration (scored as resistant; Table 2). Categorical agreement was achieved when isolates were scored as susceptible or resistant at the EUCAST breakpoint values shown in Table 1 by both methods. There was a 96% (127/132 isolate/antibiotic combinations) categorical agreement across all antibiotics tested and 100% categorical agreement for all isolates tested against trimethoprim, nitrofurantoin, cephalexin, and amoxicillin (Table 2). The microcapillary BMD had 2.3% (3/132) very major errors where 3 isolates were categorised as susceptible in contrast to microplate BMD that categorised them as resistant. All three major errors occurred with cefoxitin for which the MIC determined by microplate BMD was within ±1 log_2_ fold change of the breakpoint, indicating the breakpoint value is close to the threshold for inhibition. In this situation the variability in resistance/susceptibility scoring is known to increase significantly, leading to higher uncertainty in AST results even with gold standard methods. Microcapillary BMD showed a further 1.5% (2/132) major errors in which isolates tested for cefoxitin and ciprofloxacin were identified as resistant but categorised as susceptible by microplate BMD.

**Table 2.** AST agreement between microplate BMD and microcapillary BMD.

| Antibiotic | Microplate BMD |  | Microcapillary BMD |  | % Essential Agreement | % Categorical Agreement |
| --- | --- | --- | --- | --- | --- | --- |
|  | Susceptible | Resistant | Susceptible | Resistant |  |  |
| Trimethoprim | 18% (4/22) | 82% (18/22) | 18% (4/22) | 82% (18/22) | **100%** (55/55) | **100%** (22/22) |
| Nitrofurantoin | 100% (22/22) | 0% (0/22) | 100% (22/22) | 0% (0/22) |  | **100%** (22/22) |
| Cephalexin | 14% (3/22) | 86% (19/22) | 14% (3/22) | 86% (19/22) |  | **100%** (22/22) |
| Amoxicillin | 5% (1/22) | 95% (21/22) | 5% (1/22) | 95% (21/22) |  | **100%** (22/22) |
| Ciprofloxacin | 18% (4/22) | 82% (18/22) | 17% (3/22) | 86% (19/22) |  | **95%** (21/22) |
| Cefoxitin | 27% (6/22) | 73% (16/22) | 36% (8/22) | 64% (14/22) |  | **82%** (18/22) |

Overall, we conclude that microcapillary BMD performs very well and that the miniaturisation of broth microdilution from microplate wells down to microcapillaries has little if any impact on MIC determination and scoring for antibiotic resistance.

### 3.2 Time-resolved growth analysis in microcapillary test strips allows earlier endpoint MIC giving accurate antibiotic susceptibility within 6h

Previously we established that even at low cell densities, *E. coli* growth can be detected within 7 h and at the inoculum recommended for AST within approximately 3-5 h (Needs et al., 2021) (Fig S1). MIC were measured and susceptibility/resistance scored for microcapillary BMD after 6 h incubation (Fig S2-7), and results compared to overnight microplate BMD results (Table 3). In only two cases, MIC was not determinable due to no growth detected at 6h in the no antibiotic control, for *E. coli* 13352 tested against trimethoprim and cefoxitin. No MIC value was attributed for 9 microcapillary nitrofurantoin BMD test strips, in which growth was only detected in the growth control without antibiotic (i.e. MIC below lowest concentration; scored as susceptible), or for a further 72 microcapillary BMD test strips where all antibiotic containing capillaries showed growth and thus the MIC was above the highest concentration (scored as resistant; Table 3).

**Table 3.**
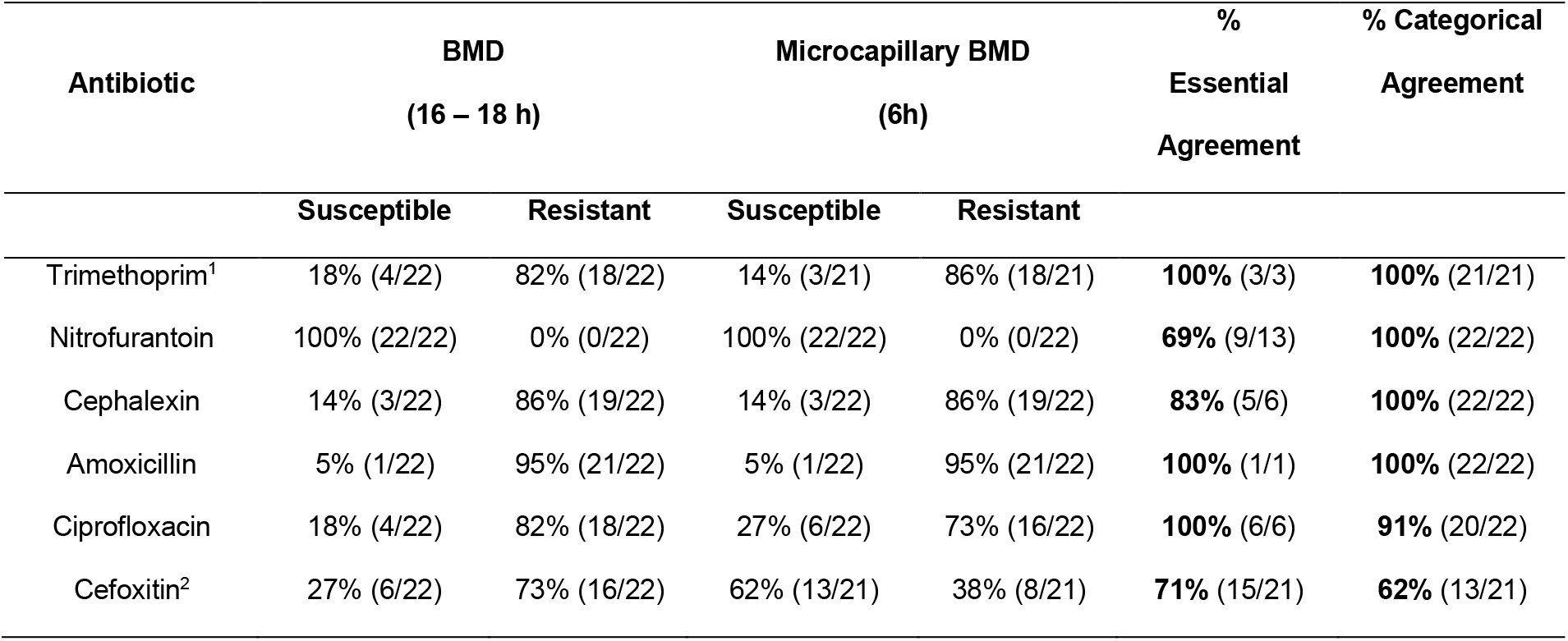
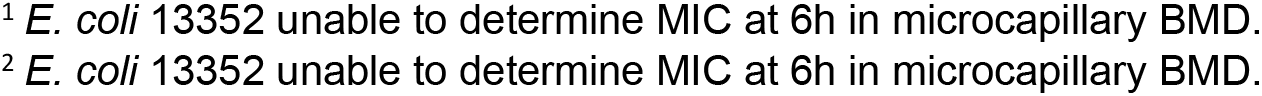
Agreement between microcapillary test strips read at 6h compared to overnight.

| Antibiotic | BMD<br>(16 – 18 h) |  | Microcapillary BMD<br>(6h) |  | %<br>Essential<br>Agreement | % Categorical<br>Agreement |
| --- | --- | --- | --- | --- | --- | --- |
|  | Susceptible | Resistant | Susceptible | Resistant |  |  |
| Trimethoprim<sup>1</sup> | 18% (4/22) | 82% (18/22) | 14% (3/21) | 86% (18/21) | **100%** (3/3) | **100%** (21/21) |
| Nitrofurantoin | 100% (22/22) | 0% (0/22) | 100% (22/22) | 0% (0/22) | **69%** (9/13) | **100%** (22/22) |
| Cephalexin | 14% (3/22) | 86% (19/22) | 14% (3/22) | 86% (19/22) | **83%** (5/6) | **100%** (22/22) |
| Amoxicillin | 5% (1/22) | 95% (21/22) | 5% (1/22) | 95% (21/22) | **100%** (1/1) | **100%** (22/22) |
| Ciprofloxacin | 18% (4/22) | 82% (18/22) | 27% (6/22) | 73% (16/22) | **100%** (6/6) | **91%** (20/22) |
| Cefoxitin<sup>2</sup> | 27% (6/22) | 73% (16/22) | 62% (13/21) | 38% (8/21) | **71%** (15/21) | **62%** (13/21) |

<sup>1</sup> *E. coli* 13352 unable to determine MIC at 6h in microcapillary BMD.
<sup>2</sup> *E. coli* 13352 unable to determine MIC at 6h in microcapillary BMD.

For those conditions where MIC was within the measurable range, overall essential agreement in MIC level across the antibiotics tested dropped to 78% (38/49 isolate/antibiotic combinations). However, overall categorical agreement remained high at 91% (120/132 isolate/antibiotic combinations). Very major errors were identified in 7.7% (10/130) of cases. The accuracy was reduced in a number of ways. Where growth was not clearly detected at the 6h timepoint, because the antibiotic slows growth, false susceptibility could be scored, but at an overnight timepoint even the slower growth still leads to a positive score. On the other hand, slower growing isolates growth may not be strongly detected at 6h even in control without antibiotic; for these strips susceptibility and MIC cannot be scored.

Kinetics of growth was clearly influenced more by some but not other antibiotics, depending on susceptibility/resistant status. Cefalexin and ciprofloxacin show very similar MIC results at 6 and 16 h varying by only one doubling dilution of antibiotic (Table S3, Fig 3a-b). Isolates categorised as resistant show indistinguishable growth kinetics in the presence of antibiotics (Figure 3c). Nitrofurantoin, in which all isolates were categorised as susceptible, showed the most distinct delay in growth kinetics in the presence of antibiotic (Fig 3d). Shorter incubation times may be less suitable for high resolution quantitation of MIC, and may be more accurate for some antibiotics than others, however overall accuracy of microcapillary BMD for scoring resistance vs sensitivity remained high even with a 6h endpoint readout. As with overnight endpoint, the growth could readily be recorded using a smartphone camera or if necessary, scored by eye. Overall, this indicates that early endpoints can be reliable for determining resistance and may aid in decision making for resistant isolates.

**Figure 3.**
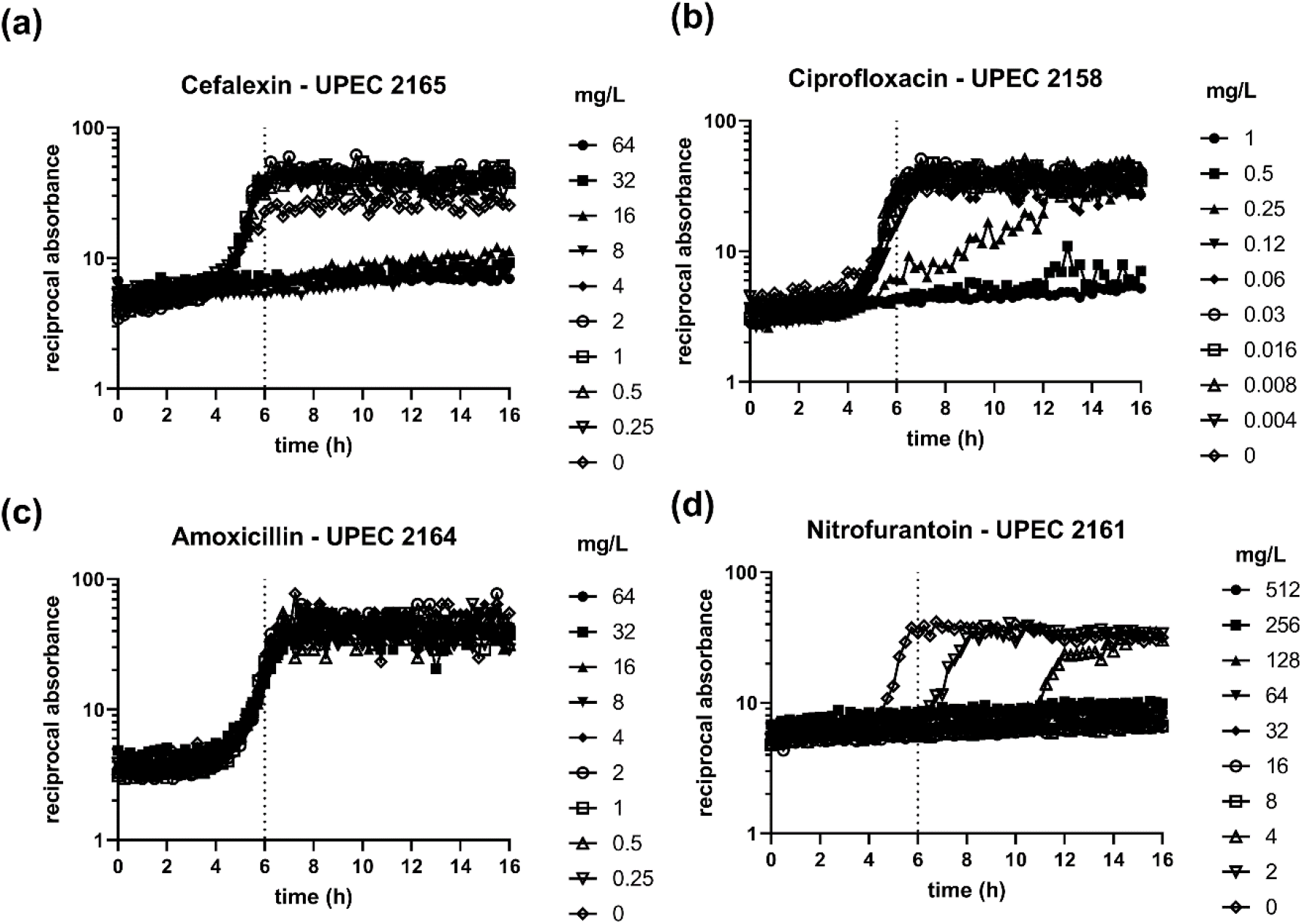
Microcapillary BMD with a fixed inoculum density growth kinetics in the presence of antibiotics. **(a)** UPEC 2165 isolate gives the same MIC result after 6h incubation compared to overnight. **(b)** UPEC 2158 recorded MIC is within 1 doubling dilution of antibiotic for ciprofloxacin. **(c)** Growth of UPEC 2164 is identical at all antibiotic concentrations of amoxicillin. **(d)** Nitrofurantoin shows the greatest amount of delayed growth. UPEC 2161 only observes a positive growth control at 6h, indicated an MIC ≤2 mg/L compared to an endpoint MIC of 8 mg/L.

### 3.3 Effect of inoculum density on microfluidic MIC determination

Detecting antibiotic susceptibility and measuring MIC is known to be significantly affected by the initial inoculum density, and in the development of new methodologies it is vital to identify how sensitive results are to variables such as inoculum cell density that are not easy to control. Here, we focussed on understanding how starting cell density affects microcapillary BMD performance, and seeing if IE reduces accuracy for both MIC quantitation and AST scoring, as it does for conventional microplate methods. Microcapillary BMD test strips were dipped into serial 10-fold dilutions of each bacterial isolate, starting with 10^7^ CFU/mL (Fig 4), 100X higher than the recommended density. Inoculum densities 100X above the CLSI recommended resulted in growth at antibiotic concentrations significantly above the MIC, giving false resistance scores for *E. coli* 25922 and UPEC 2165 for ciprofloxacin and cefalexin and in UPEC 2165 for nitrofurantoin (Fig 4b and Table 4). This confirms that as for microplate BMD, the starting inoculum should be standardised to avoid false resistance from high starting cell density.

**Figure 4.**
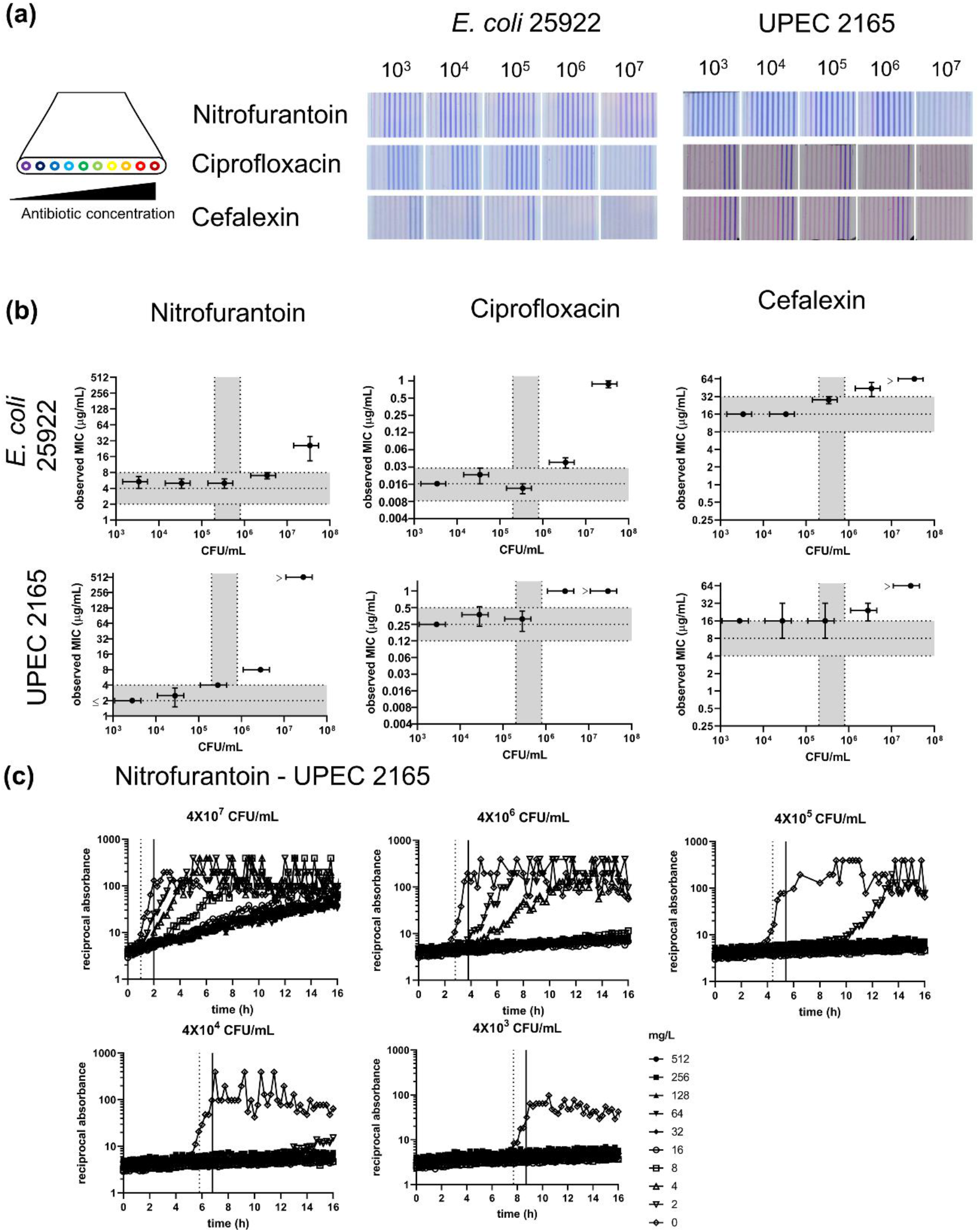
Endpoint MIC measurement of isolates using range of inoculum density. **(a)** Schematic showing antibiotic cocnetration is highest in capilary 10. Capillary 1 contains no antibtioc and acts as a growth control. Representative images of capillary test strips at various incoculum density ranging from 10^3^ – 10^7^ CFU/mL. **(b)** MIC observed at different inoculum density. Data indicates average of duplicate capillaries for two independent experiments. Vertical error bars indicate ± SEM and horizontal error bars indicate ± SD for inoclum density determiend by overnight agar plating. **(c)** Growth kinetics of UPEC 2165 against nitrofurantoin. Dotted line indicates detection time for the growth control, solid line indicates 1 h cuttoff from growth control. Data points indicate average of duplicate capillaries.

**Table 4.**
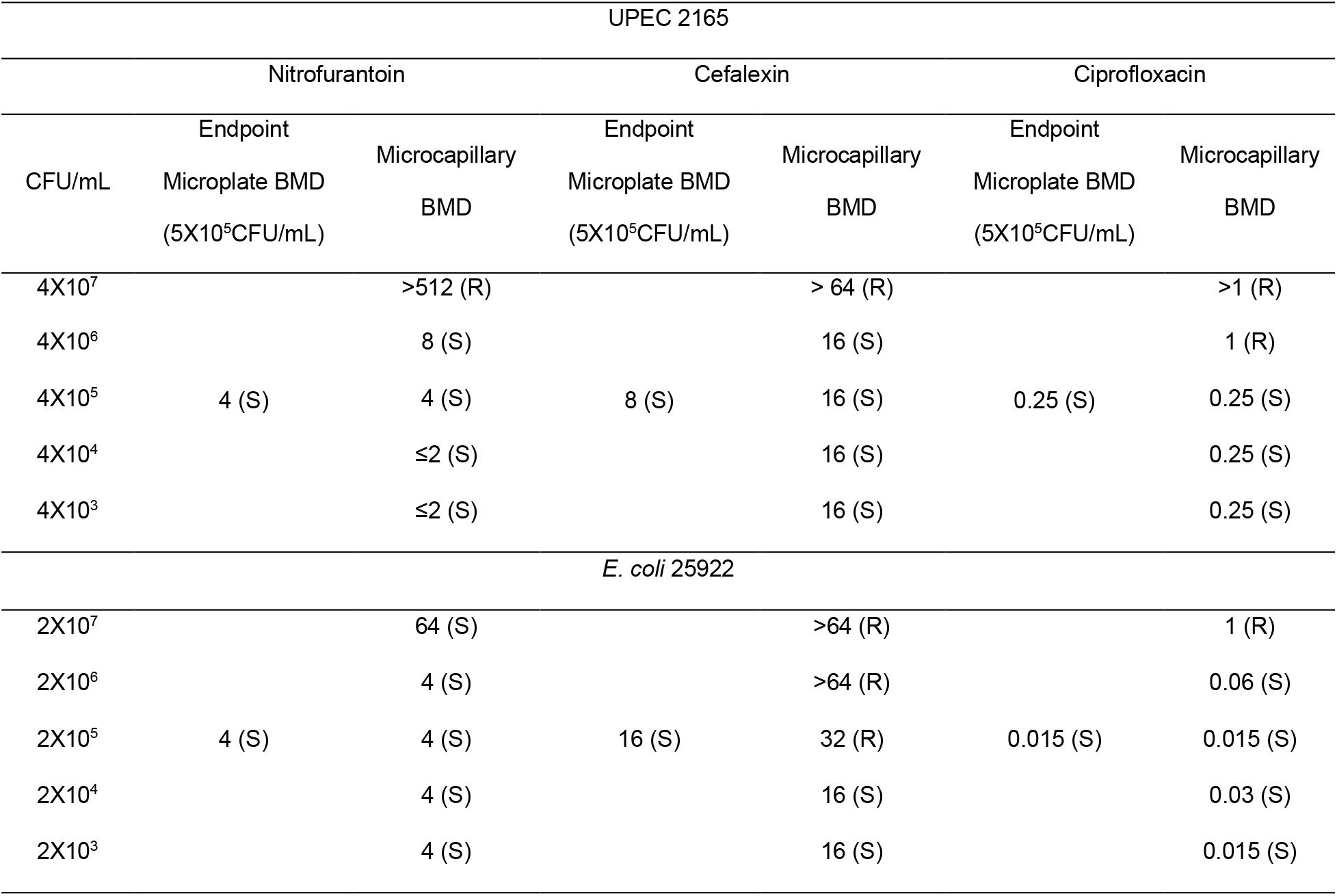
Determination of endpoint MIC using varying inoculum density.

Inoculum densities below the CLSI recommended density gave MIC measurements within the ± 1 log_2_ fold acceptable range, suggesting the microcapillary BMD is relatively robust as long as inoculum density remains low enough to avoid false resistance or raised MIC levels. This indicates that the microcapillary BMD test strips are compatible with lower bacterial inoculum, suggesting this method may be robust enough to be used in environments where precise inoculum density may be harder to achieve (e.g. less well-resourced laboratories).

### 3.4 Detecting delay in growth from kinetic analysis allows fast and accurate categorical determination at higher cell densities

Rapid AST microdevices have been developed that use time-lapse digital microscopy imaging to determine delayed growth in the presence of antibiotics compared to a growth control (Avesar et al., 2017, Kang et al., 2019, Busche et al., 2019, Sun et al., 2019, Osaid et al., 2021). Higher inoculum densities convert resazurin faster. The use of miniaturised BMD tests combined with kinetic data collection offers further opportunity to gain more detailed insight into bacterial response to antimicrobial agents, for example through detection of differential growth kinetics indicating susceptibility. MIC was calculated based on differential growth kinetics for different inoculum densities (Table 5, Fig S8-9). MIC was calculated by scoring susceptibility when growth was delayed, but resistant when at a particular antibiotic concentration, growth was detected within 1 h of the no antibiotic control. It was evident that isolates that were categorised as susceptible, and with MIC well below the breakpoint concentration, were consistently scored as susceptible by this method, even at inoculum densities 100X higher than the recommended range. Nitrofurantoin showed significantly delayed growth compared to the growth control at all inoculum densities (Fig 4c). False resistance at higher inocula was still observed for some isolate/antibiotic combinations but with an MIC still within one doubling dilution of the breakpoint antibiotic concentration. For example, *E. coli* 25922 has an MIC for ciprofloxacin of 0.015 μg/mL. Based on growth kinetics, inocula as high as 10^7^ CFU/mL would be scored as susceptible. The UPEC isolate 2165 with an MIC of 0.25 μg/mL – the same value as the breakpoint – exhibited false resistance at inoculum densities greater than 10^5^ CFU/mL. It is clear therefore that the fastest way to score resistance or susceptibility, or to estimate MIC, is through detecting delayed growth with a high inoculum density that allows early growth detection. The accuracy of this method varies depending on the antibiotic, and as with even gold standard methods, if the MIC of isolates is close to breakpoint concentrations, AST is less reliable.

**Table 5.** MIC determination based on delayed growth in the presence of antibiotics.

| UPEC 2165 |  |  |  |  |  |  |  |
| --- | --- | --- | --- | --- | --- | --- | --- |
|  |  | Nitrofurantoin |  | Cefalexin |  | Ciprofloxacin |  |
| CFU/mL | Growth Control -<br>Time to<br>resazurin<br>conversion<br>(h) | Endpoint<br>Microplate BMD<br>(5X10<sup>5</sup>CFU/mL) | Kinetic<br>MIC | Endpoint<br>Microplate BMD<br>(5X10<sup>5</sup>CFU/mL) | Kinetic<br>MIC | Endpoint<br>Microplate BMD<br>(5X10<sup>5</sup>CFU/mL) | Kinetic MIC |
| 4X10<sup>7</sup> | 1 |  | 8 (S) |  | > 64 (R) |  | >1 (R) |
| 4X10<sup>6</sup> | 2.8 |  | ≤2 (S) |  | 16 (S) |  | 1 (R) |
| 4X10<sup>5</sup> | 4.4 | 4 (S) | ≤2 (S) | 8 (S) | 16 (S) | 0.25 (S) | 0.25 (S) |
| 4X10<sup>4</sup> | 5.8 |  | ≤2 (S) |  | 16 (S) |  | 0.25 (S) |
| 4X10<sup>3</sup> | 7.7 |  | ≤2 (S) |  | 8 (S) |  | 0.125 (S) |
| *E. coli* 25922 |  |  |  |  |  |  |  |
| 2X10<sup>7</sup> | 1.5 |  | ≤2 (S) |  | >64 (R) |  | 0.25 (S) |
| 2X10<sup>6</sup> | 3.1 |  | ≤2 (S) |  | >64 (R) |  | 0.015 (S) |
| 2X10<sup>5</sup> | 4.9 | 4 (S) | ≤2 (S) | 16 (S) | 8 (S) | 0.015 (S) | 0.015 (S) |
| 2X10<sup>4</sup> | 5.6 |  | ≤2 (S) |  | 8 (S) |  | 0.007 (S) |
| 2X10<sup>3</sup> | 7.7 |  | ≤2 (S) |  | 4 (S) |  | 0.007 (S) |

## 4. Discussion

Overall, by measuring growth with large panels of antibiotic combined with a set of *E. coli* strains with a range of sensitivities, we observed that microfluidic broth microdilution performs very similarly to microplate broth microdilution. While agreement between the methods was very close (Table 2), the conditions where different results were obtained represented samples with marginal MIC where variability in AST is known to be highest. We observed the IE, that is well characterised for microplate measurements, also affects microfluidic broth microdilution (Fig 4), with implications for direct testing methods where bacterial inoculum density may be unknown or difficult to control. As might be expected, faster results can be obtained with earlier growth readouts, yet some trade-off is apparent between speed and accuracy. Nevertheless, even with 6h endpoint (Table 3) or when differential growth is used to score susceptibility (Table 5), accuracy remained excellent for some antibiotics and overall was 91%.

Antibiotics were chosen based on their use in UTI treatment but also to cover a range of antibiotic classes. UTIs are the most common outpatient infection (Medina and Castillo-Pino, 2019). In the UK, the 2018 NICE guidelines recommend a first-choice treatment for uncomplicated UTI in women over 16 years old of Nitrofurantoin or Trimethoprim, despite resistance rates to trimethoprim around 30% (Watts et al., 2020). During pregnancy, the need for safe and effective drugs is increased, while nitrofurantoin remains first choice, amoxicillin is only recommended if the susceptibility is known due to resistance rates above 50% in some demographics (Watts et al., 2020). Cefoxitin, while not currently used for clinical prescription, has been cited as an effective *in vitro* against UTI pathogens including extended-spectrum beta-lactamase producing Enterobacteriaceae (ESBL-E) (Mambie et al., 2016, Senard et al., 2018, Guet-Revillet et al., 2014). Since antibiotic loading of the strips is by freeze-drying, it is expected that the test can be adapted to most antibiotics for different applications.

Bacterial growth was monitored using the metabolic indicator resazurin. Resazurin has been used in many different plate assays studying bacterial and cellular growth, mitochondrial function, and MIC of antimicrobial agents. However, with many cell viability test methods the resazurin is added towards the end of the experiment and incubated for several hours to measure metabolic activity after a primary incubation period with antimicrobial agent but no dye, to reduce the risk of the dye interacting with the antimicrobial (Kim and Jang, 2018, Teh et al., 2017, Rakhmawatie et al., 2019, Elshikh et al., 2016, Sarker et al., 2007). In contrast, for the novel microcapillary BMD method, the resazurin is present from the start. Microplate BMD with the addition of resazurin at the start was therefore compared to microplate BMD without dye to evaluate if the presence of resazurin dye affected antimicrobial susceptibility and MIC results. There was 100% essential agreement when microplate BMD was compared with and without resazurin dye for the MIC values of nitrofurantoin and cephalexin, and 100% essential agreement between microcapillary BMD (that includes resazurin) vs microplate BMD conducted without resazurin (Table S2). This confirms that the inclusion of resazurin does not interfere with MIC determination for *E. coli* when mixed with the sample at 0.25 mg/mL and incubated overnight. Resazurin can be used with fluorescence detection (Needs et al., 2021) with the potential for earlier growth detection compared to colorimetric detection, however, in previous studies, some isolates showed a decay in fluorescence after the initial increase (Needs et al., 2019). This has been observed previously in assays in which fluorescent resorufin is further reduced to the colorless, non -fluorescence hydroresorufin. This reduces the reliability of an endpoint measurement as loss of fluorescent signal may be mistakenly interpreted as negative growth i.e. scored as false susceptible (Uzarski et al., 2017). For this reason, we measured growth colorimetrically. Although small, the individual capillaries are large enough to be scored by the naked eye if digital camera is not available to record result. A smartphone camera is ideal for digitally recording these test results, and wide range of different quality digital cameras have been shown to be equally effective for recording microcapillary bioassay results (Needs et al., 2021, Jégouic et al., 2021).

The ability to detect growth of very low viable cell numbers – ideally down to a single colony forming unit – may be critical for some analytical applications. We previously identified the limit of detection of a 1 μL microcapillary test strip to be in the order of 5×10^3^ CFU/mL; at this concentration 99% of capillaries are predicted to contain at least 1 bacteria every test, allowing MIC measurement (Needs et al., 2021). The initial inoculum density correlates to the speed of detecting resazurin conversion, as early growth at low density does not convert enough dye to influence the capillary color, thus lower densities take longer before growth can be detected (Needs et al., 2021). Thus, although reduced inoculum density still gave categorical agreement with microplate BMD, early timepoint endpoint readings at 6h are not possible as growth is not always detected without antibiotic at this timepoint (Fig S1).

Whilst overnight plating to obtain individual colony isolates, plus cell density adjustment prior to testing is still required, these results confirm that microcapillary BMD has equivalent performance to standard microplate BMD. The MCF test strips significantly increase throughput compared to microplates alone, whilst remaining compatible with conventional microtitre plate format found in most laboratories for sample preparation. While one standard microplate BMD can test 6 isolates in duplicate with 6 concentrations of a single antibiotic, the microcapillary BMD can test 96 isolates, with a single isolate per well. Each microcapillary BMD test strip provides 9 antibiotic concentrations plus control, and each well can be dipped with multiple test strips for replicates. Assuming picking and adjusting isolate concentration, and plating samples out, is rate limiting, microcapillary BMD is therefore 16 times higher throughput than microplates.

## 4 Conclusions

We present a high throughput method for determining antibiotic resistance or susceptibility and for measuring MIC, using a simple to use microfluidic dipstick device, and validate this method with uropathogenic *E. coli* and antibiotics commonly used to treat UTI. The test is fully compatible with existing lab equipment such as multichannel pipettes and 96-well format microplates for sample preparation and does not require additional equipment such as pumps or controllers for microfluidic liquid handling. The colorimetric detection of bacterial growth allows easy readout and simple analysis, reducing the need for equipment investment – most digital cameras including smartphone cameras are capable of recording color change, or growth can be scored by eye. The microcapillary BMD can provide rapid categorical susceptibility/resistance scoring at time points within 6 h based on differential growth. Kinetic analysis can provide robust S/R categorisation above the recommended inoculum density, however, isolates with MIC within two-fold dilution of the breakpoint may show false resistance.

## Author Contributions

SHN: Conceptualisation, Data curation, Formal analysis, Investigation, Methodology, Project administration, Software, Visualization, Writing – original draft, Writing – review & editing. WI: Resources, Writing – review & editing. ZR: Resources, Writing – review & editing. SA: Resources, Writing – review & editing. PR: Writing – review & editing. ADE: Conceptualisation, Funding acquisition, Methodology, Project administration, Supervision, Writing – original draft, Writing – review & editing

## Acknowledgements

Figure 1a was adapted from elements from Servier medical art under CC BY 3.0 licence.

## Ethical Approval

The collection of urinary pathogenic *E. coli* from a tertiary care hospital of Pakistan from community acquired UTI patients and was approved by Ethical Review Board (ERB) of Pakistan Institute of Medical Sciences.

