## Supporting Information for "High-throughput, multiplex microfluidic test strip for the determination of antibiotic susceptibility in uropathogenic *E. coli* with smartphone detection"

### S1.1 MIC values provided for microplate BMD and microcapillary BMD tests

MIC values are provided for all experiments summarized in the main text. Values provided represent the highest MIC value of at least duplicate wells/capillaries unless otherwise specified.

**Table S1. Minimum inhibitory concentration in uropathogenic *E. coli* isolates in microtiter plate (BMD) and microcapillary BMD (MCF).**

| | | MIC (mg/L) | | | | | | | | | | | |
|---|---|---|---|---|---|---|---|---|---|---|---|---|---|
| | | Cefoxitin | | Ciprofloxacin | | Trimethoprim | | Nitrofurantoin | | Cephalexin | | Amoxicillin | |
| | | BMD | MCF | BMD | MCF | BMD | MCF | BMD | MCF | BMD | MCF | BMD | MCF |
| Ref. Strain | ATCC 25922 | 4 | 8 | 0.015 | 0.015 | 0.5 | 1 | 4 | 8 | 16 | 16 | 4 | 8 |
| | NCTC 13352 | 4 | 2 | 0.015 | 0.03 | 0.5 | 0.25 | ≤2 | 8 | 16 | 8 | >64 | >64 |
| UPEC Isolate ID | 2151 | 8 | 8 | >1 | >1 | >32 | >32 | 8 | 8 | >64 | >64 | >64 | >64 |
| | 2152 | 16 | 16 | >1 | >1 | >32 | >32 | 4 | 8 | >64 | >64 | >64 | >64 |
| | 2153 | 32 | 16 | >1 | >1 | >32 | >32 | 8 | 8 | 64 | 32 | >64 | >64 |
| | 2154 | 16 | 8 | >1 | >1 | >32 | >32 | 8 | 8 | >64 | >64 | >64 | >64 |
| | 2155 | 16 | 8 | >1 | >1 | >32 | >32 | 8 | 8 | 32 | 64 | >64 | >64 |
| | 2156 | 32 | 16 | 0.5 | 0.5 | >32 | >32 | 16 | 16 | >64 | >64 | >64 | >64 |
| | 2157 | 32 | 32 | >1 | >1 | >32 | >32 | 8 | 8 | 64 | 64 | >64 | >64 |
| | 2158 | 16 | 16 | 0.25 | 0.5 | >32 | >32 | 16 | 8 | >64 | >64 | >64 | >64 |
| | 2159 | 16 | 8 | >1 | >1 | >32 | >32 | 16 | 16 | >64 | >64 | >64 | >64 |
| | 2160 | 32 | 64 | >1 | >1 | >32 | >32 | 8 | 8 | >64 | >64 | >64 | >64 |
| | 2161 | 16 | 16 | >1 | >1 | >32 | >32 | 4 | 8 | >64 | >64 | >64 | >64 |
| | 2162 | 32 | 32 | >1 | >1 | >32 | >32 | 8 | 16 | >64 | >64 | >64 | >64 |
| | 2163 | 16 | 16 | 0.5 | 1 | >32 | >32 | 4 | 4 | >64 | >64 | >64 | >64 |
| | 2164 | 8 | 8 | >1 | >1 | >32 | >32 | 8 | 16 | >64 | >64 | >64 | >64 |
| | 2165 | 4 | 8 | 0.25 | 0.25 | >32 | >32 | ≤2 | 4 | 8 | 8 | >64 | >64 |
| | 2166 | 16 | 16 | >1 | >1 | 2 | 2 | 16 | 16 | >64 | >64 | >64 | >64 |
| | 2167 | 32 | 64 | >1 | >1 | >32 | >32 | 8 | 8 | >64 | >64 | >64 | >64 |
| | 2168 | 64 | 32 | >1 | >1 | >32 | >32 | 4 | 8 | >64 | >64 | >64 | >64 |
| | 2169 | 8 | 16 | >1 | >1 | >32 | >32 | ≤2 | 4 | >64 | >64 | >64 | >64 |
| | 2170 | 64 | 32 | >1 | >1 | 0.5 | 0.25 | 64 | 64 | >64 | >64 | >64 | >64 |

**Table S2. Comparison between MIC determined by microplate BMD with and without resazurin and microcapillary BMD with resazurin.**

| | | MIC (µg/mL) | | | | | |
|---|---|---|---|---|---|---|---|
| | | **Nitrofurantoin** | | | **Cephalexin** | | |
| | | **Microplate BMD** | **Microplate BMD + resazurin** | **Microcapillary BMD + resazurin** | **Microplate BMD** | **Microplate BMD + resazurin** | **Microcapillary BMD + resazurin** |
| **Ref Strain** | ATCC 25922 | 4 | 4 | 8 | **16** | 16 | 16 |
| | NCTC 13352 | <2 | <2 | 8 | **16** | 16 | 8 |
| **UPEC Isolate ID** | 2151 | 8 | 8 | 8 | **>64** | >64 | >64 |
| | 2152 | 8 | 4 | 8 | **>64** | >64 | >64 |
| | 2153 | 8 | 8 | 8 | **32** | 64 | 32 |
| | 2154 | 8 | 8 | 8 | **>64** | >64 | >64 |
| | 2155 | 8 | 8 | 8 | **32** | 32 | 64 |
| | 2156 | 16 | 16 | 16 | >64 | >64 | >64 |
| | 2157 | 16 | 8 | 8 | **64** | 64 | 64 |
| | 2158 | 16 | 16 | 8 | >64 | >64 | >64 |
| | 2159 | 16 | 16 | 16 | >64 | >64 | >64 |
| | 2160 | 8 | 8 | 8 | >64 | >64 | >64 |
| | 2161 | 8 | 4 | 8 | >64 | >64 | >64 |
| | 2162 | 16 | 8 | 16 | >64 | >64 | >64 |
| | 2163 | 4 | 4 | 4 | >64 | >64 | >64 |
| | 2164 | 8 | 8 | 16 | >64 | >64 | >64 |
| | 2165 | 4 | ≤2 | 4 | **8** | 8 | 8 |
| | 2166 | 16 | 16 | 16 | >64 | >64 | >64 |
| | 2167 | 8 | 4 | 8 | >64 | >64 | >64 |
| | 2168 | 8 | 4 | 8 | >64 | >64 | >64 |
| | 2169 | 4 | ≤2 | 4 | >64 | >64 | >64 |
| | 2170 | 128 | 64 | 64 | >64 | >64 | >64 |

**Table S3. MIC determined by microplate BMD (BMD) after 16 h incubation and MIC determined by microcapillary BMD (MCF) after 6 h incubation.**

| | | Cefoxitin | | Ciprofloxacin | | Trimethoprim | | Nitrofurantoin | | Cephalexin | | Amoxicillin | |
|---|---|---|---|---|---|---|---|---|---|---|---|---|---|
| | | **BMD** | **MCF (6h)** | **BMD** | **MCF (6h)** | **BMD** | **MCF (6h)** | **BMD** | **MCF (6)** | **BMD** | **MCF (6)** | **BMD** | **MCF (6)** |
| **Ref. Strain** | **ATCC 25922** | 4 | 2 | 0.015 | 0.015 | 0.5 | 1 | 4 | 4 | 16 | 8 | 4 | 4 |
| | **NCTC 13352** | 4 | N.D. | 0.015 | 0.007 | 0.5 | N.D. | ≤2 | ≤2 | 16 | 4 | >64 | >64 |
| **UPEC Isolate ID** | **2151** | 8 | 4 | >1 | >1 | >32 | >32 | 8 | 4 | >64 | >64 | >64 | >64 |
| | **2152** | 16 | 4 | >1 | >1 | >32 | >32 | 4 | ≤2 | >64 | >64 | >64 | >64 |
| | **2153** | 32 | 16 | >1 | >1 | >32 | >32 | 8 | 4 | 64 | 64 | >64 | >64 |
| | **2154** | 16 | 4 | >1 | >1 | >32 | >32 | 8 | ≤2 | >64 | >64 | >64 | >64 |
| | **2155** | 16 | 8 | >1 | >1 | >32 | >32 | 8 | 4 | 32 | 32 | >64 | >64 |
| | **2156** | 32 | 8 | 0.5 | 0.25 | >32 | >32 | 16 | 4 | >64 | >64 | >64 | >64 |
| | **2157** | 32 | 32 | >1 | >1 | >32 | >32 | 8 | ≤2 | 64 | 64 | >64 | >64 |
| | **2158** | 16 | 16 | 0.25 | 0.25 | >32 | >32 | 16 | 4 | >64 | >64 | >64 | >64 |
| | **2159** | 16 | 4 | >1 | >1 | >32 | >32 | 16 | 4 | >64 | >64 | >64 | >64 |
| | **2160** | 32 | 32 | >1 | >1 | >32 | >32 | 8 | 4 | >64 | >64 | >64 | >64 |
| | **2161** | 16 | 8 | >1 | >1 | >32 | >32 | 4 | ≤2 | >64 | >64 | >64 | >64 |
| | **2162** | 32 | 32 | >1 | >1 | >32 | >32 | 8 | 4 | >64 | >64 | >64 | >64 |
| | **2163** | 16 | 4 | 0.5 | 0.25 | >32 | >32 | 4 | ≤2 | >64 | >64 | >64 | >64 |
| | **2164** | 8 | 8 | >1 | >1 | >32 | >32 | 8 | 4 | >64 | >64 | >64 | >64 |
| | **2165** | 4 | 4 | 0.25 | 0.25 | >32 | >32 | ≤2 | ≤2 | 8 | 8 | >64 | >64 |
| | **2166** | 16 | 4 | >1 | >1 | 2 | 1 | 16 | 4 | >64 | >64 | >64 | >64 |
| | **2167** | 32 | 16 | >1 | >1 | >32 | >32 | 8 | ≤2 | >64 | >64 | >64 | >64 |
| | **2168** | 64 | 32 | >1 | >1 | >32 | >32 | 4 | 4 | >64 | >64 | >64 | >64 |
| | **2169** | 8 | 8 | >1 | >1 | >32 | >32 | ≤2 | ≤2 | >64 | >64 | >64 | >64 |
| | **2170** | 64 | 32 | >1 | >1 | 0.5 | 0.25 | 64 | 32 | >64 | >64 | >64 | >64 |

### S1.2 Growth kinetics of *E. coli* grown in microcapillary film

(a)

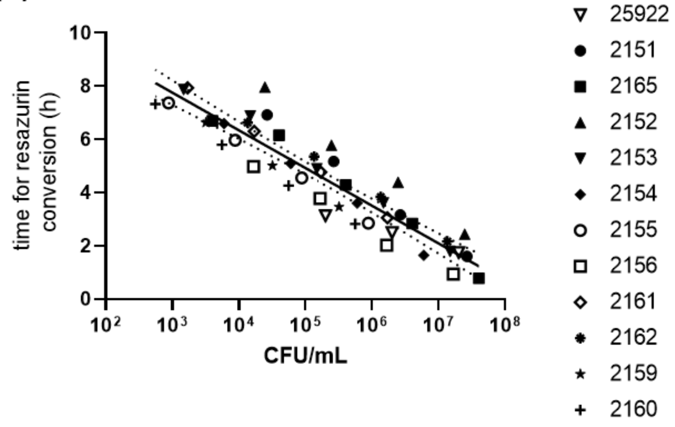


(b)

| Time (h) | | | |
|---|---|---|---|
| | Mean | Upper 95% CI | Lower 95% CI |
| $2X10^5$ CFU/mL | 4.5 | 4.7 | 4.3 |
| $8X10^5$ CFU/mL | 3.6 | 3.9 | 3.3 |

**Figure S1. Time to resazurin conversion is plotted against CFU/ml determined by overnight spot dot.** Shaded area indicates CLSI standard inoculum range for MTP BMD. Solid line indicates linear regression and dotted lines indicate 95% confidence intervals **(e)** Table indicates the average time to growth detection at two different inoculum cell densities, and 95% confidence intervals for detection time. Data points indicate the average of 3-10 capillaries

### S1.3 Growth kinetics of *E. coli* grown in the presence of antibiotics

Growth curves for all isolates tested are provided. Graphs indicate the reciprocal absorbance in the red channel.

Cefoxitin

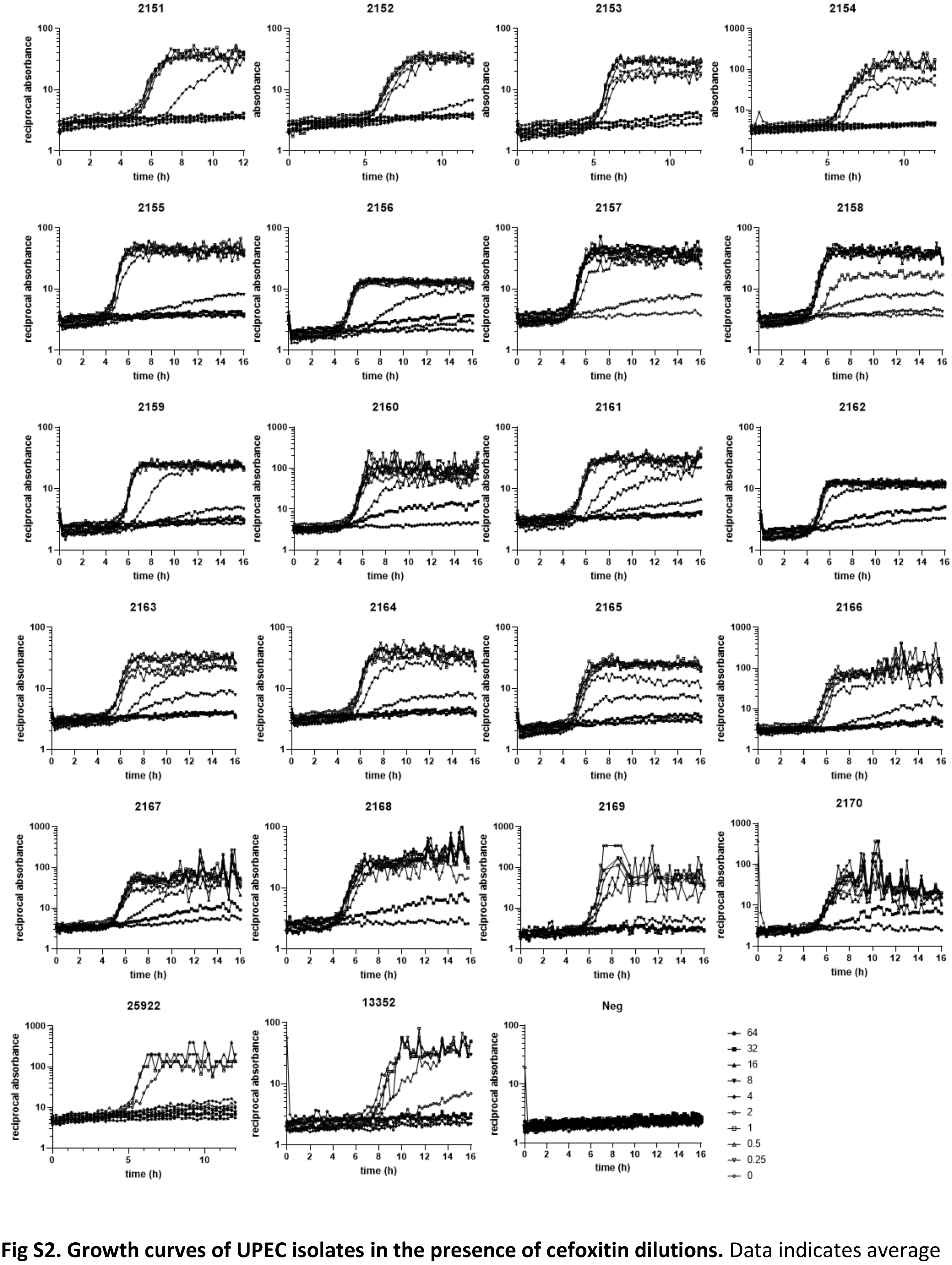


**Fig S2. Growth curves of UPEC isolates in the presence of cefoxitin dilutions.** Data indicates average of duplicate capillaries.

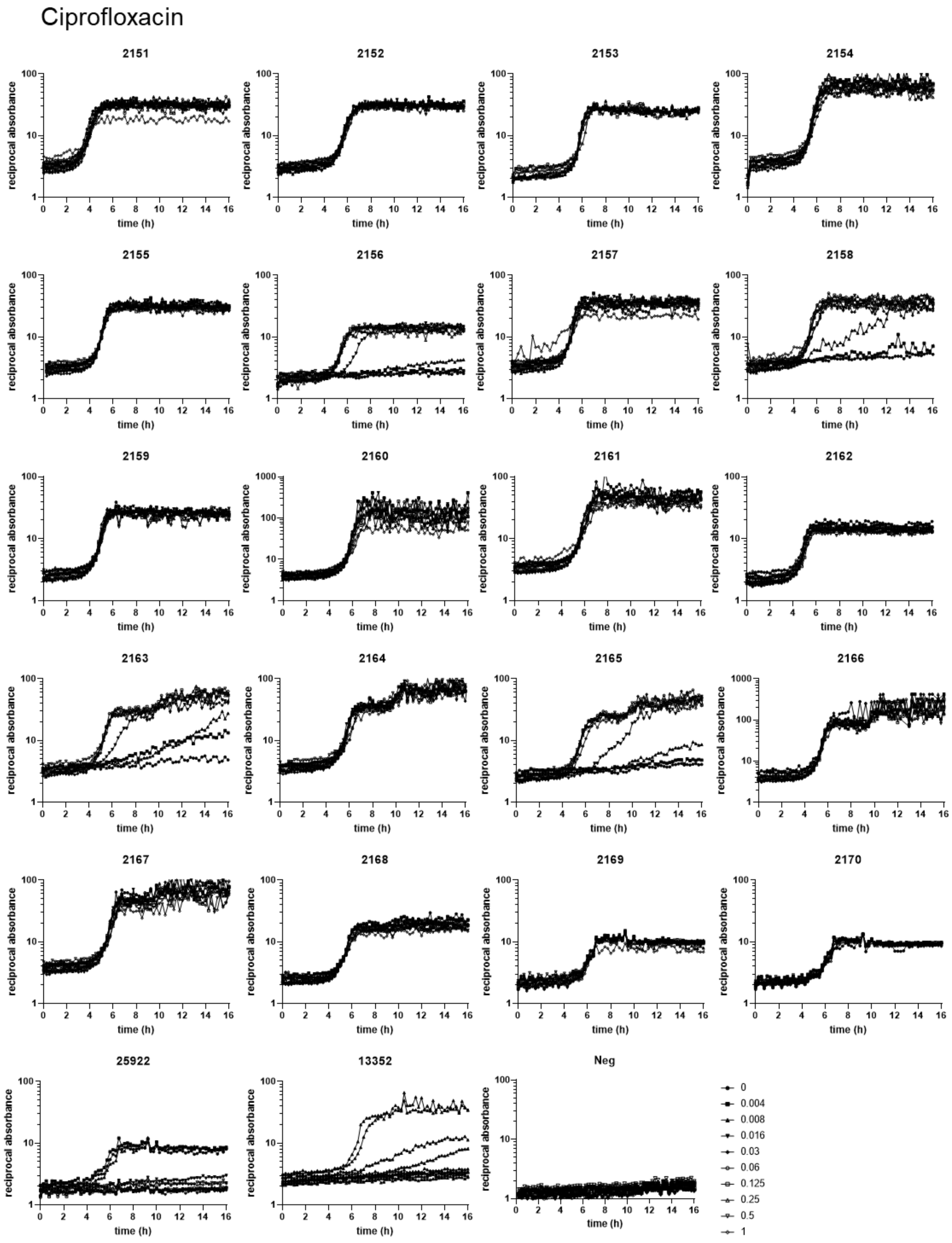


**Fig S3. Growth curves of UPEC isolates in the presence of ciprofloxacin dilutions.** Data indicates average of duplicate capillaries.

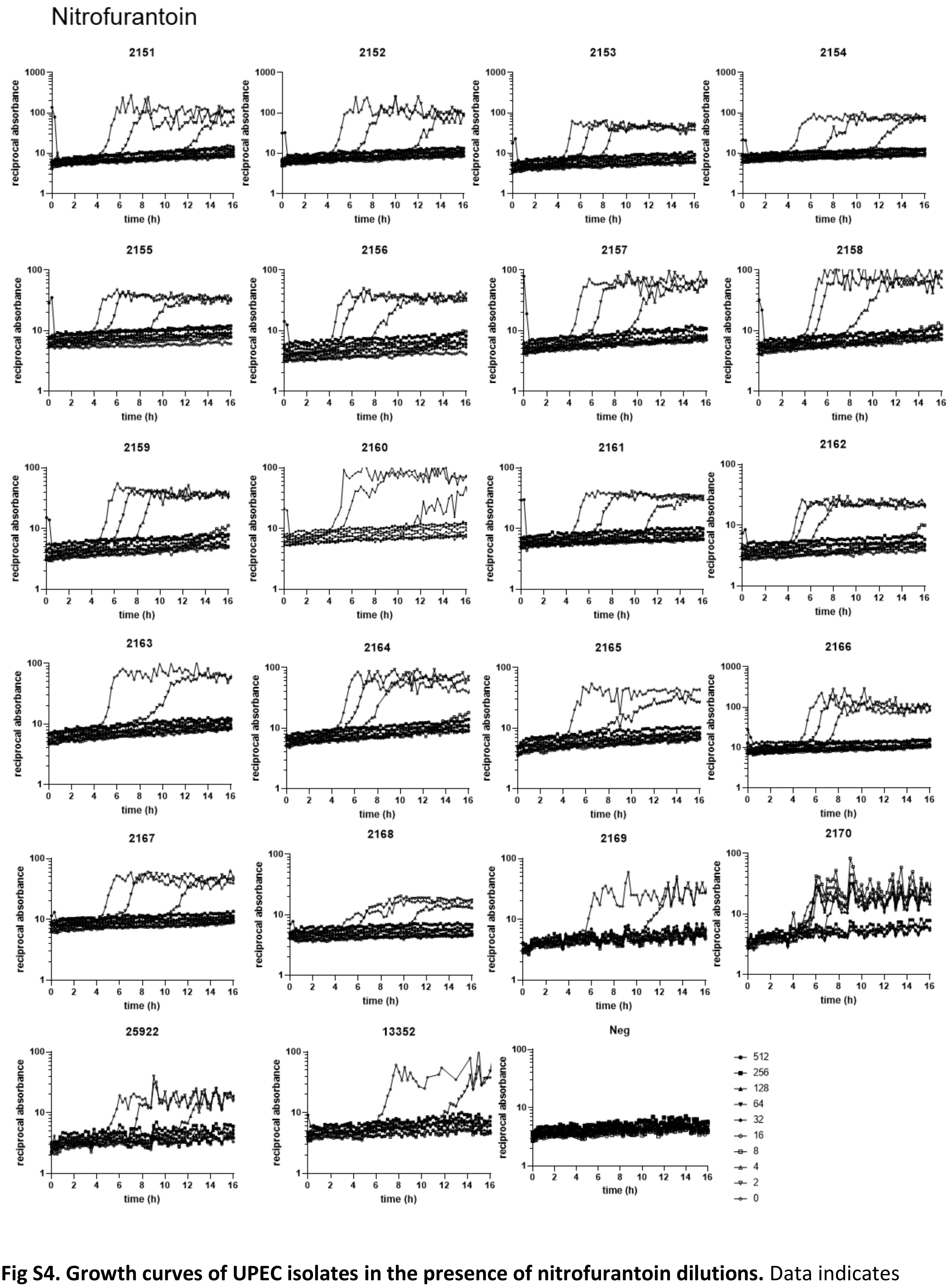


**Fig S4. Growth curves of UPEC isolates in the presence of nitrofurantoin dilutions.** Data indicates average of duplicate capillaries.

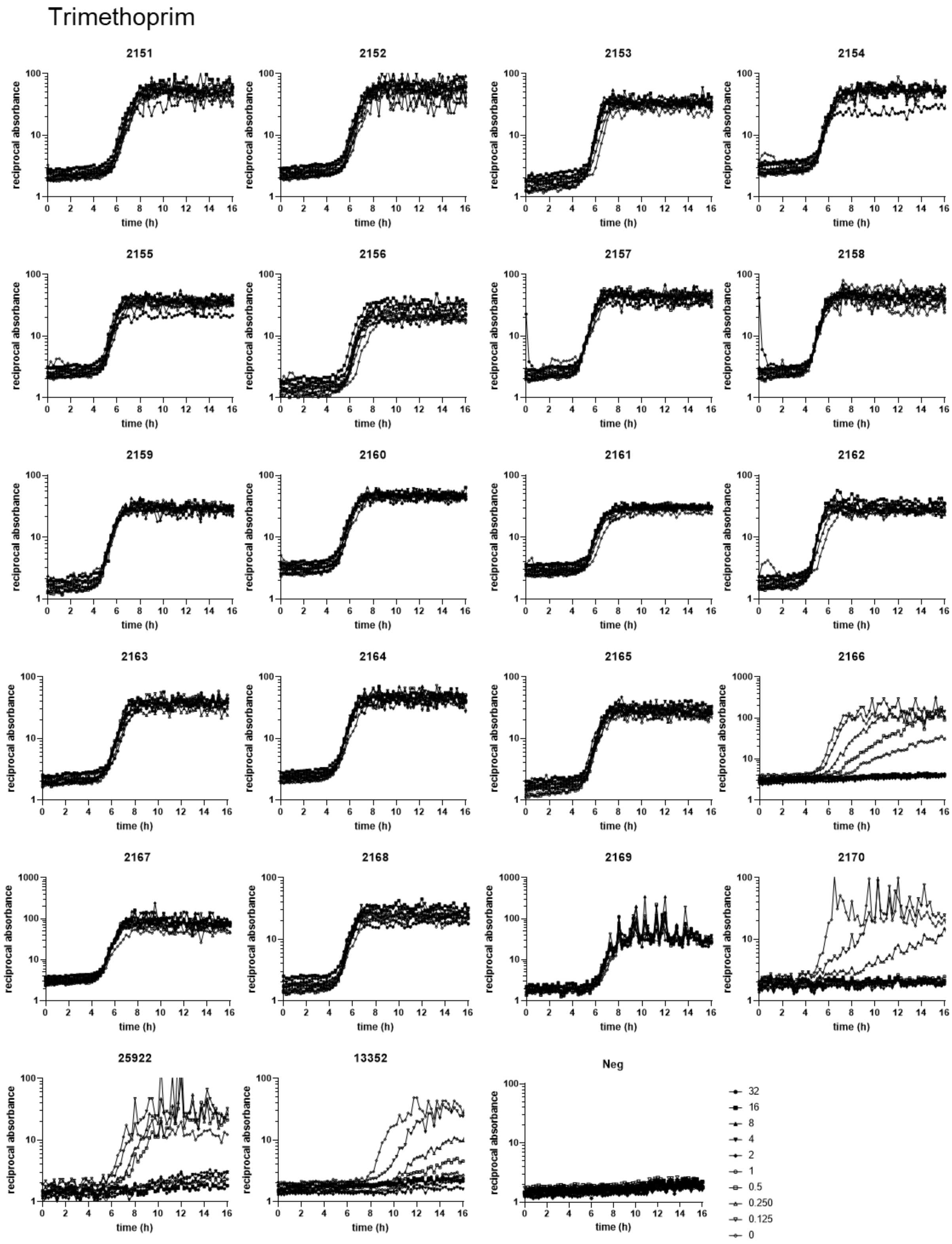


**Fig S5. Growth curves of UPEC isolates in the presence of trimethoprim dilutions.** Data indicates average of duplicate capillaries.

Cefalexin

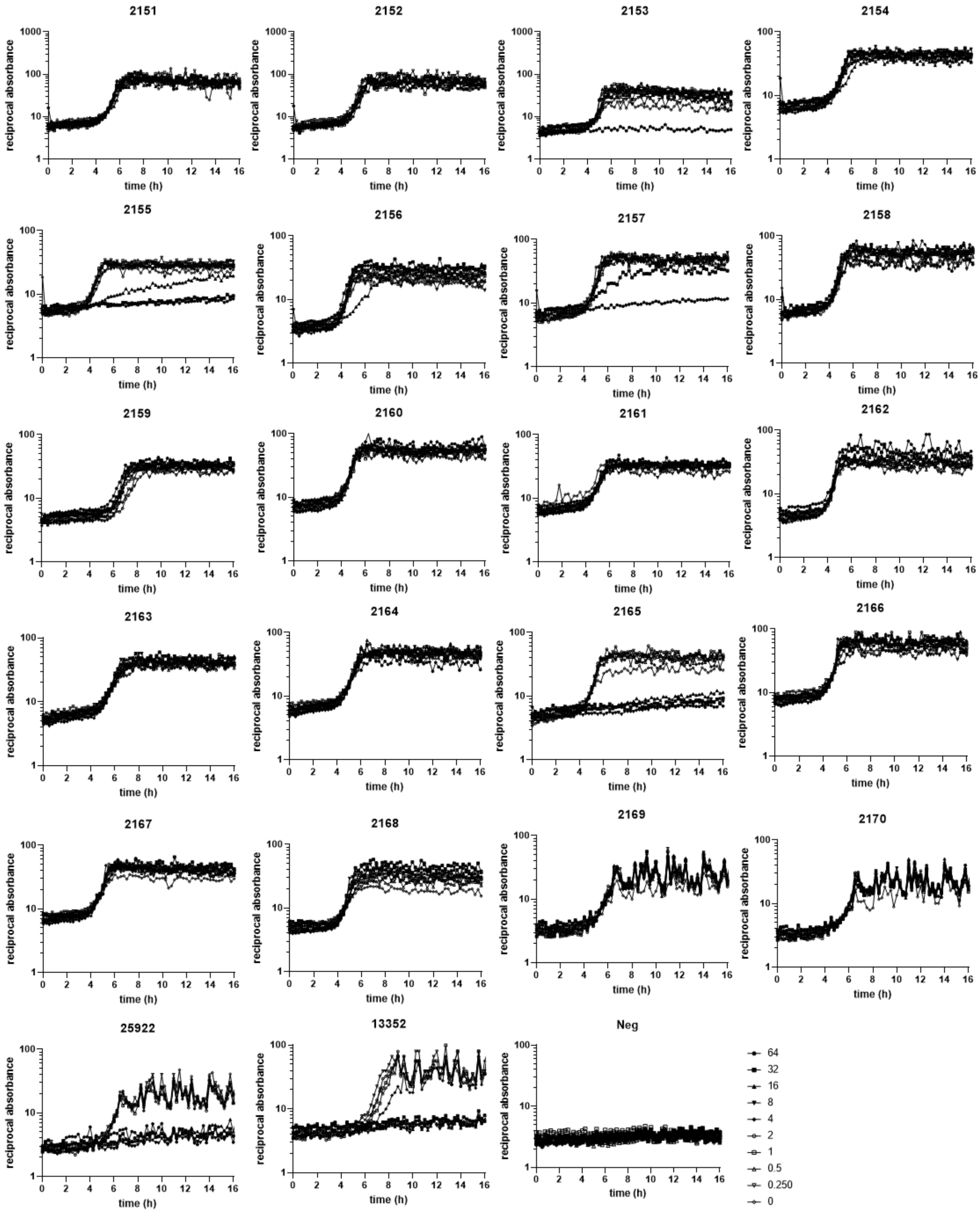


**Fig S6. Growth curves of UPEC isolates in the presence of cefalexin dilutions.** Data indicates average of duplicate capillaries.

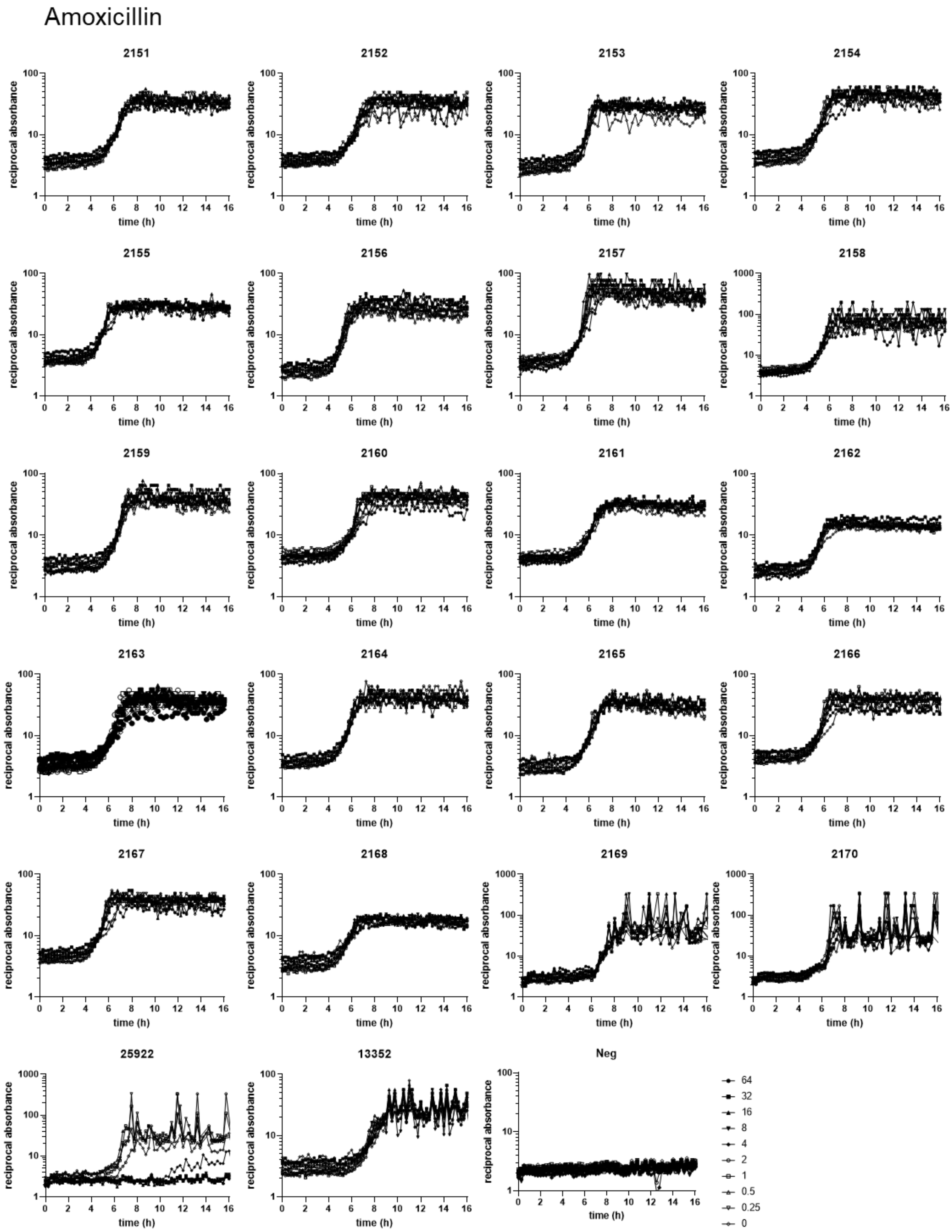


**Fig S7. Growth curves of UPEC isolates in the presence of amoxicillin dilutions.** Data indicates average of duplicate capillaries.

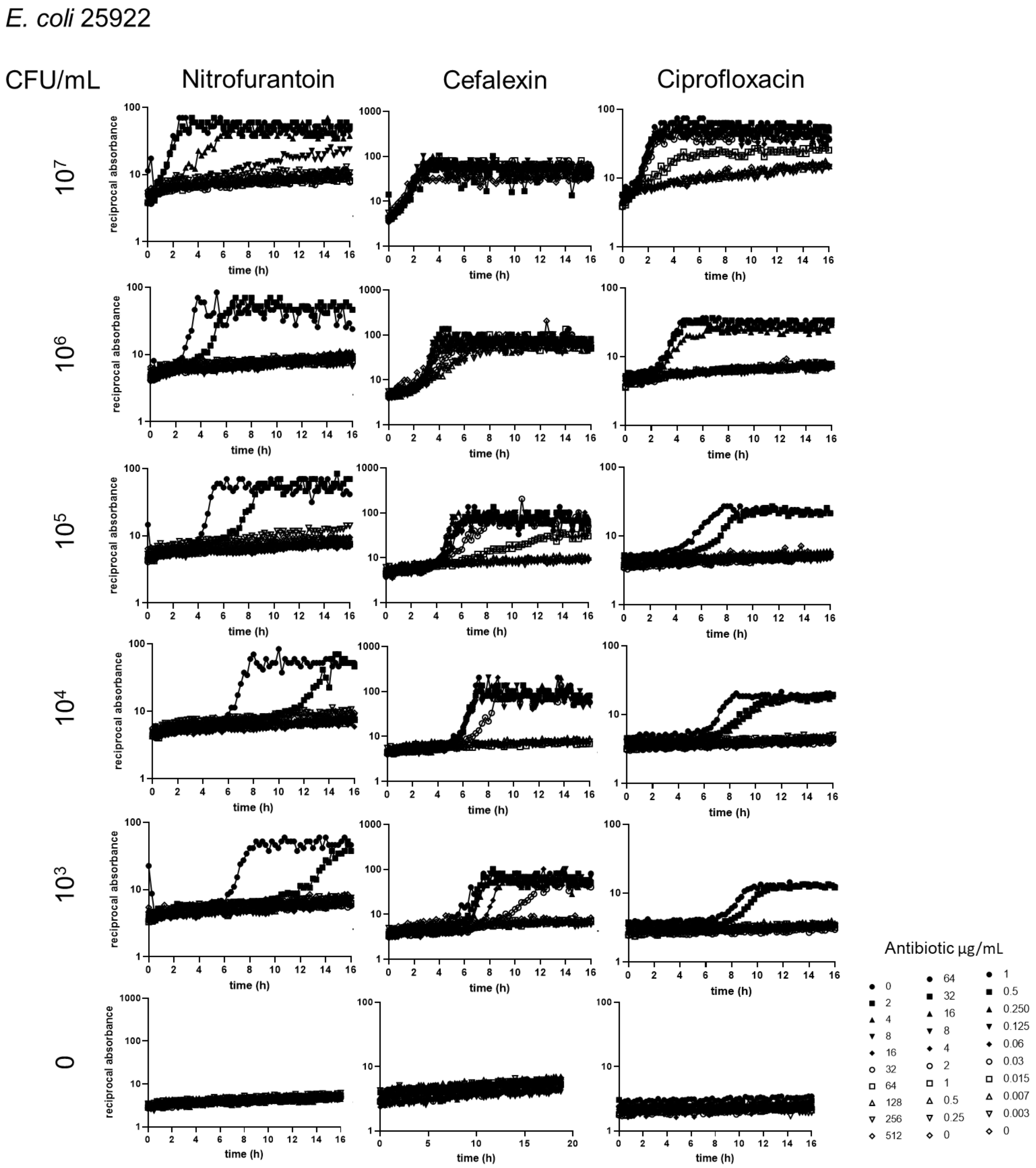


**Figure S8. Growth curves of *E. coli* 25922 at different starting inoculum densities in the presence of dilutions of nitrofurantoin, cefalexin and ciprofloxacin.** Data indicates average of duplicate capillaries.

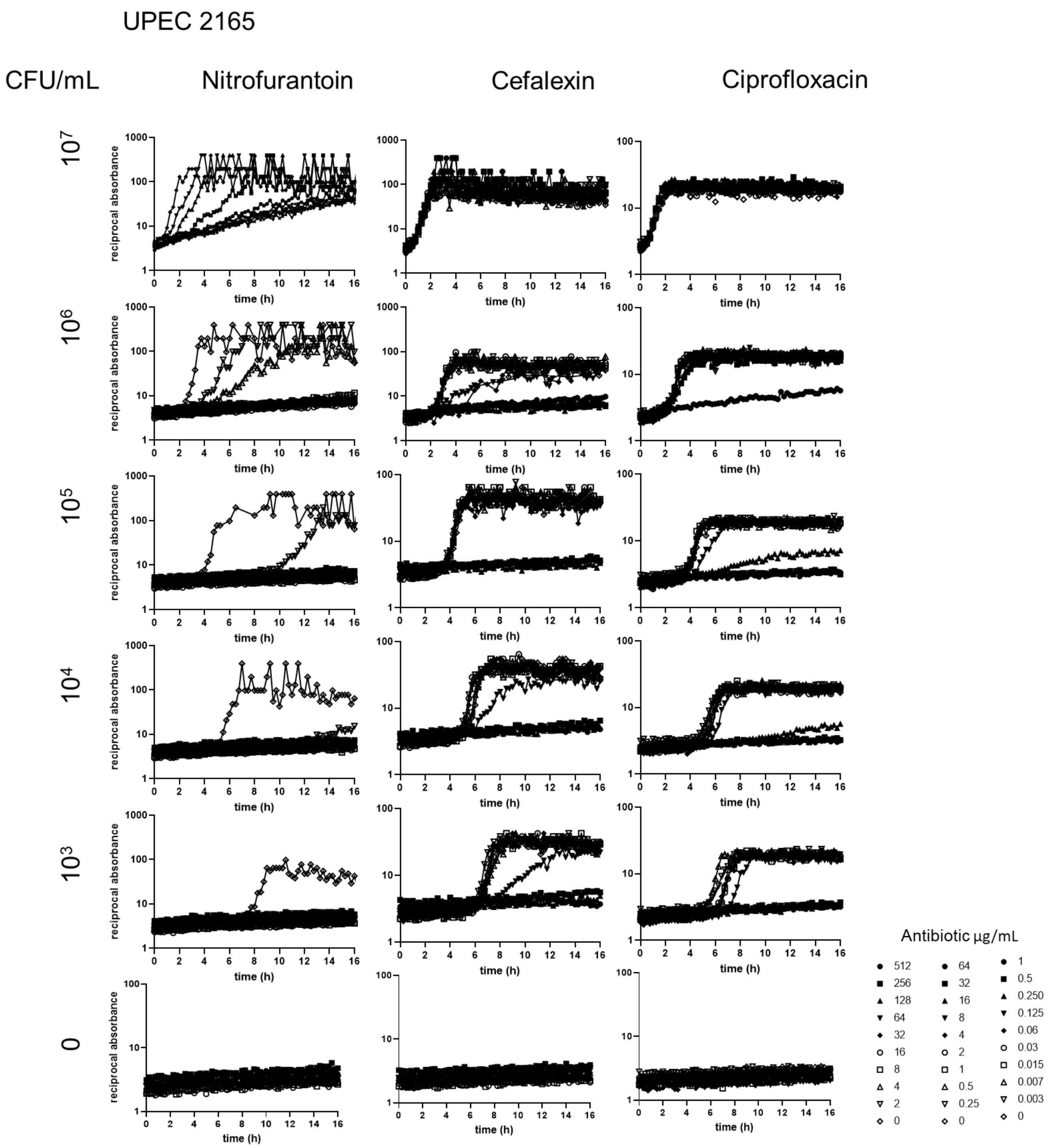


**Figure S9. Growth curves of UPEC 2165 at different starting inoculum densities in the presence of dilutions of nitrofurantoin, cefalexin and ciprofloxacin.** Data indicates average of duplicate capillaries.
